# Infant gut strain persistence is associated with maternal origin, phylogeny, and functional potential including surface adhesion and iron acquisition

**DOI:** 10.1101/2021.01.26.428340

**Authors:** Yue Clare Lou, Matthew R. Olm, Spencer Diamond, Alexander Crits-Christoph, Brian A. Firek, Robyn Baker, Michael J. Morowitz, Jillian F. Banfield

**Affiliations:** Department of Plant and Microbial Biology, University of California, Berkeley, CA 94720, USA; Department of Microbiology and Immunology, Stanford University School of Medicine, Stanford, CA 94305, USA; Department of Earth and Planetary Science, University of California, Berkeley, CA 94709, USA; Department of Surgery, University of Pittsburgh School of Medicine, Pittsburgh, PA 15213, USA; Department of Environmental Science, Policy, and Management, University of California, Berkeley, CA 94720, USA; Earth Sciences Division, Lawrence Berkeley National Laboratory, Berkeley, CA 94705, USA; Chan Zuckerberg Biohub, San Francisco, CA 94158, USA

## Abstract

Gut microbiome succession impacts infant development. However, it remains unclear what factors promote persistence of initial bacterial colonists in the developing gut. Here, we performed strain-resolved metagenomic analyses to compare gut colonization of preterm and full-term infants throughout the first year of life and evaluated links between strain persistence and strain origin as well as genetic potential. Analysis of 206 fecal metagenomes collected from full-term and preterm infants and their mothers revealed that infants’ initially distinct microbial communities converged by age one. Approximately 11% of early colonists, primarily *Bacteroides* and *Bifidobacterium*, persisted during the first year of life, and these were more prevalent in full-term compared to preterm infants. Gut-associated strains from mothers were significantly more likely to persist in the infant gut than other strains. Enrichment in genes for surface adhesion, iron acquisition and carbohydrate degradation may explain persistence of some strains through the first year of life.

## INTRODUCTION

Microorganisms rapidly colonize the near sterile infant gut during and shortly after birth (Robertson et al., 2019). These early gut colonizers play important roles in the maturation of infants’ metabolic pathways, especially related to the immune system (Wang et al., 2020). Disruptions of early-life microbial acquisition and assembly via factors such as antibiotic treatment have been associated with increased risk of developing diseases later in life including asthma, allergies, and metabolic syndrome (Arrieta et al., 2015; Bisgaard et al., 2011; Tamburini et al., 2016). Certain bacterial commensals can persist within the adult gut for years (Faith et al., 2013; Schloissnig et al., 2013; Zhao et al., 2019). Infant gut microbiomes are less stable than adult microbiomes at the whole-community level (Koenig et al., 2011; Yatsunenko et al., 2012), and fundamental questions remain regarding the persistence of their early colonists. There is potential for long-term impact if the first colonizing strains, which are often hospital-associated pathogens in premature infants (Gasparrini et al., 2019; Gibson et al., 2016; Raveh-Sadka et al., 2016), persist as infants develop. Thus, it is important to analyze strain persistence and the sources and characteristics of persisting strains, as well as the time required for convergence of premature and full-term microbiomes.

The majority of studies on the infant microbiome have relied on 16S rRNA sequencing, which cannot resolve genomic differences beyond the species level. These studies have advanced our understanding of the early-life gut microbiota assembly process (Koenig et al., 2011; La Rosa et al., 2014; Yatsunenko et al., 2012). However, in order to answer questions regarding organism transmissions from various sources, organism persistence through early life, or sharing of organisms amongst individuals, whole genome resolution is necessary. Robust detection of subtle genomic differences allows one to determine whether strains are identical or merely closely related, and to distinguish commensal from pathogenic strains (Brito and Alm, 2016; Olm et al., 2021). Sequencing cultured isolates is one way to recover microbial genomes, but it is low throughput, targeted to particular taxa, and unlikely to capture the full strain diversity present (Van Rossum et al., 2020). Genome-resolved metagenomics circumvents the shortcomings of 16S rRNA and culture-dependent sequencing by rapidly generating genomes for essentially all microorganisms present in the gastrointestinal tracts of infants early in life without relying on culturing or any public reference genomes (Brooks et al., 2014, 2017; Olm et al., 2017a, 2019a).

Here, we investigated early-life gut microbiome assembly dynamics using genome-resolved metagenomics. Our study targeted preterm and full-term infants born at the same hospital over a three-year period, and tracked their gut microbiome compositions to age one. We also collected fecal samples from mothers at birth to identify transmission of strains between the infant and maternal gut microbiomes. Our work is distinct in that we examined the succession of the early-life gut microbiomes with rigorous strain-level resolution, which allowed us to accurately track the fate and gene content of strains colonizing the infant intestinal tract. Taken together, we determined that maternal origin, phylogeny and functional potential of bacterial early colonists all contributed to early-life strain persistence in infants. Insights regardings traits that enable gut microbiome residency during early-life have implications for development of rational microbiome manipulations.

## RESULTS

### Study design and sampling

In this study, we followed 23 full-term and 19 preterm infants from birth to age one. A total of 402 fecal metagenomes from these infants and their mothers were selected and subjected to deep metagenomic sequencing (~3.5 Tbp of total sequence data in the form of 150 bp paired-end reads) (Figure S1). Reads were *de novo* assembled to recover 7521 draft genome bins, which were further dereplicated at 98% whole-genome average nucleotide identity (gANI) to yield 1,005 genomes that represent unique microbial “subspecies”. We use the term subspecies as a taxonomic rank in between strain and species (Figure 1) (Methods).

**Figure 1.**
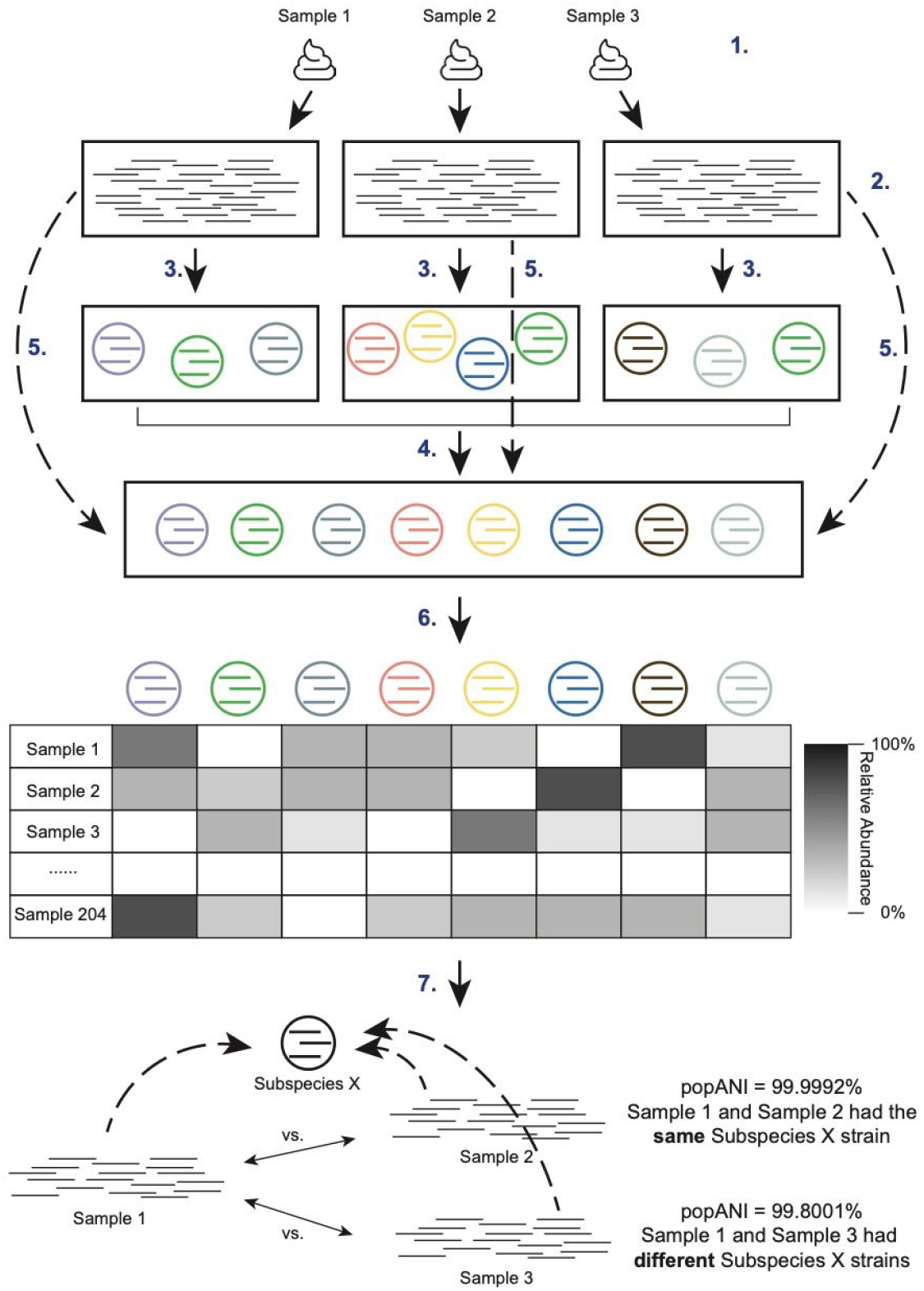
Genome-resolved metagenomics pipeline. Fecal samples collected from infants and their mothers (1) were subjected to shotgun sequencing (2). Reads were subsequently processed and assembled into draft genomes (3), which were dereplicated together at 98% gANI to result in 1,005 genomes that represent unique microbial subspecies (4). Reads from each individual sample were then mapped to all 1005 subspecies (5) for sample-specific genome detection (detection threshold = 1x coverage across ≥50% of the genome) and sample-specific relative abundance calculation (6). inStrain was run to identify same strains of the subspecies between samples of the same or different infants. Specifically, inStrain was run to compare the genome similarity among all subspecies that were present in ≥2 samples (7).

Detection of identical strains was achieved using inStrain (Olm et al., 2021) and was based on comparisons of read mapping to the same subspecies. A bacterial bin was considered identical in two samples if the compared region of the genome from both samples shared over 99.999% population-level ANI (popANI) based on previously suggested thresholds (Olm et al., 2021) (Figure 1). Our stringent definition of “strain” allowed us to discriminate between recent strain transmission events and pairs of organisms that shared a recent evolutionary history but originated from distinct sources. The use of high resolution genome-enabled population metagenomic comparisons distinguishes our work from that of prior studies that used public reference genomes and substantially less stringent strain definitions (Ferretti et al., 2018; Korpela et al., 2018; Yassour et al., 2018).

In addition to infant and maternal samples, we sequenced five negative reagent controls (one per extraction plate). The detection of common gut species in two negative control wells prompted us to thoroughly assess artifactual sequence sharing among the wells of all extraction plates. We concluded that the contamination observed in two negative control wells was a result of well-to-well contamination on these two extraction plates (Methods). Given the importance of strain-level analyses, we rejected all samples on those plates (samples from 10 full-term and 10 preterm infants and 12 samples from mothers). No contamination was found in the other three extraction plates. Therefore, the 206 samples on these three plates (22 infants and their mothers) were used for downstream analyses (Figure S1). Metadata (Table S1) and sequencing data (Table S2) of these 22 infants and their mothers are provided.

### Approximately 11% of bacterial early colonizers persisted throughout the first year of life

Infant fecal samples were grouped into seven windows of time (months 0, 1, 2, 3, 4, 8 and 12) based on infants’ chronological ages at the time of sample collection. Bacterial strains that arrived during the first two months of life were classified as early colonists, and they were further subdivided into “persisters” or “non-persisters” depending on whether they stayed within the infant gut beyond month 8 (persisters) or not (non-persisters) using the 99.999% popANI strain identity cutoff (Methods) (Figure 2A).

**Figure 2.**
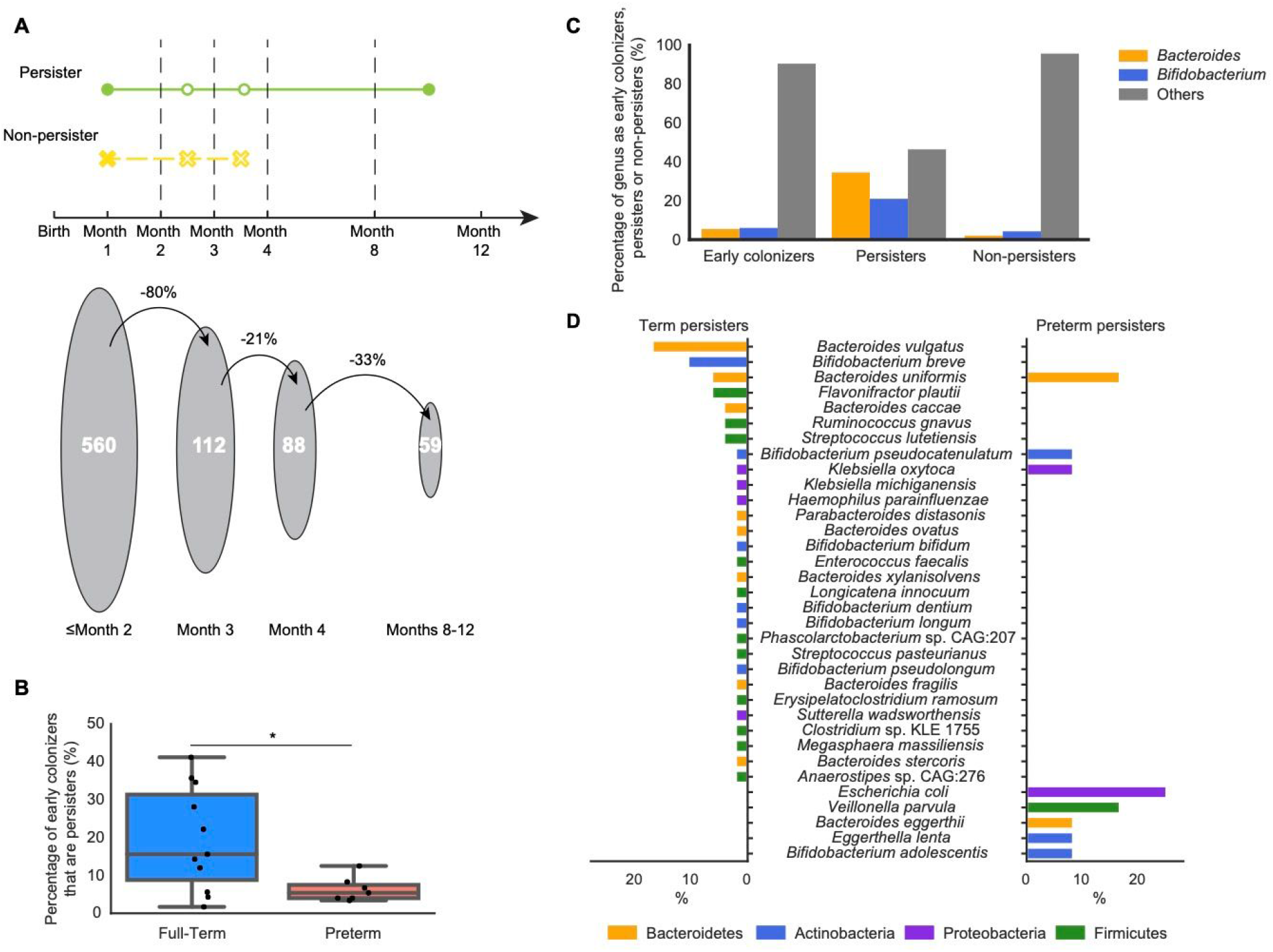
Certain bacterial strains persist in the infant gut from birth until near age one. (A) (Top) Definition of persister (present both before month 2 and after month 8) and non-persister (present before month 2 and absent after month 4) bacterial strains. (Bottom) The decrease in the number of early colonists (strains detected in the first 2 months of life) present at sequential time points. (B) Percentage of early colonizers that become persisters in the gut microbiomes of full-term and preterm infants (* = *P* < 0.05; Wilcoxon rank-sums test). (C) Percentage of early colonists, persisters and non-persisters by genus. Genera other than *Bacteroides* and *Bifidobacterium* are grouped into “Others”. (D) Species composition of persisters in full-term (left) and preterm (right) infants. Bars are colored by phylum. The x-axis is the percentage of the specific species in full-term (left) and preterm (right) persisters.

We found that 274 (47.7%) of the 575 bacterial subspecies detected across infants during the first year of life were early colonists. These 274 subspecies comprise 560 distinct strains, of which, 59 were persisters and 501 were non-persisters (Figure 2A). The median residence time for persisters was 9.6 months (95% confidence interval, 9.0–10.1 months), and the median residence time for non-persisters was 0.4 months (95% confidence interval, 0.3–0.5 months). Of the non-persisters, ~70% were not detected after month 2.

*Bacteroides* and *Bifidobacterium* strains were more likely to persist than strains of other bacterial genera (*P* = 1.1e-14 and 1.2e-04, respectively; Fisher’s exact test with FDR correction) (Figure 2B). In addition, infants who were breastfed-only before the introduction of solid foods had more persisters than those who had a mixed diet of breast milk and formula (*P* = 0.048, Wilcoxon rank-sum test). No infants in our study were formula-fed only before weaning. Notably, no Bacteroidetes persisters were found in infants who had a mixed diet, suggesting that components in breast milk such as human milk oligosaccharides (HMOs) might select for Bacteroidetes in the infant gut microbiome.

To test if prematurity had an impact on strain persistence, we compared persisters detected from full-term and preterm infants. A higher percentage of early colonizers persisted throughout the first year of life in full-term infants than in preterm infants (*P* = 0.03, Wilcoxon rank-sum test) (Figure 2C). Additionally, *B. vulgatus* and *B. breve* were more likely to persist in full-term infants (*P* = 6.0e-06 and 0.0011, respectively; Fisher’s exact test with FDR correction) whereas preterm infants had more *B. uniformis* and *E. coli* persisters (*P* = 0.016 and 0.022, respectively; Fisher’s exact test with FDR correction) (Figure 2D).

### Maternally-derived strains are more likely to be persisters in the infant gut microbiome

To elucidate the influence of maternally-derived intestinal strains on the development of the infant gut microbiome, we measured strain sharing between infants and their mothers. In this study, vertical transmission only refers to bacterial strains being transmitted from the gut microbiomes of mothers to infants, as no samples from other body sites were collected. Among the 17 infants for whom we had maternal fecal samples, 9 of the 12 full-term and 3 of the 5 preterm infants inherited strains from their mothers. In total, there were 50 maternally-sourced bacterial strains across the infant cohort (4.4% of all identified maternal strains; Figure 3A).

**Figure 3.**
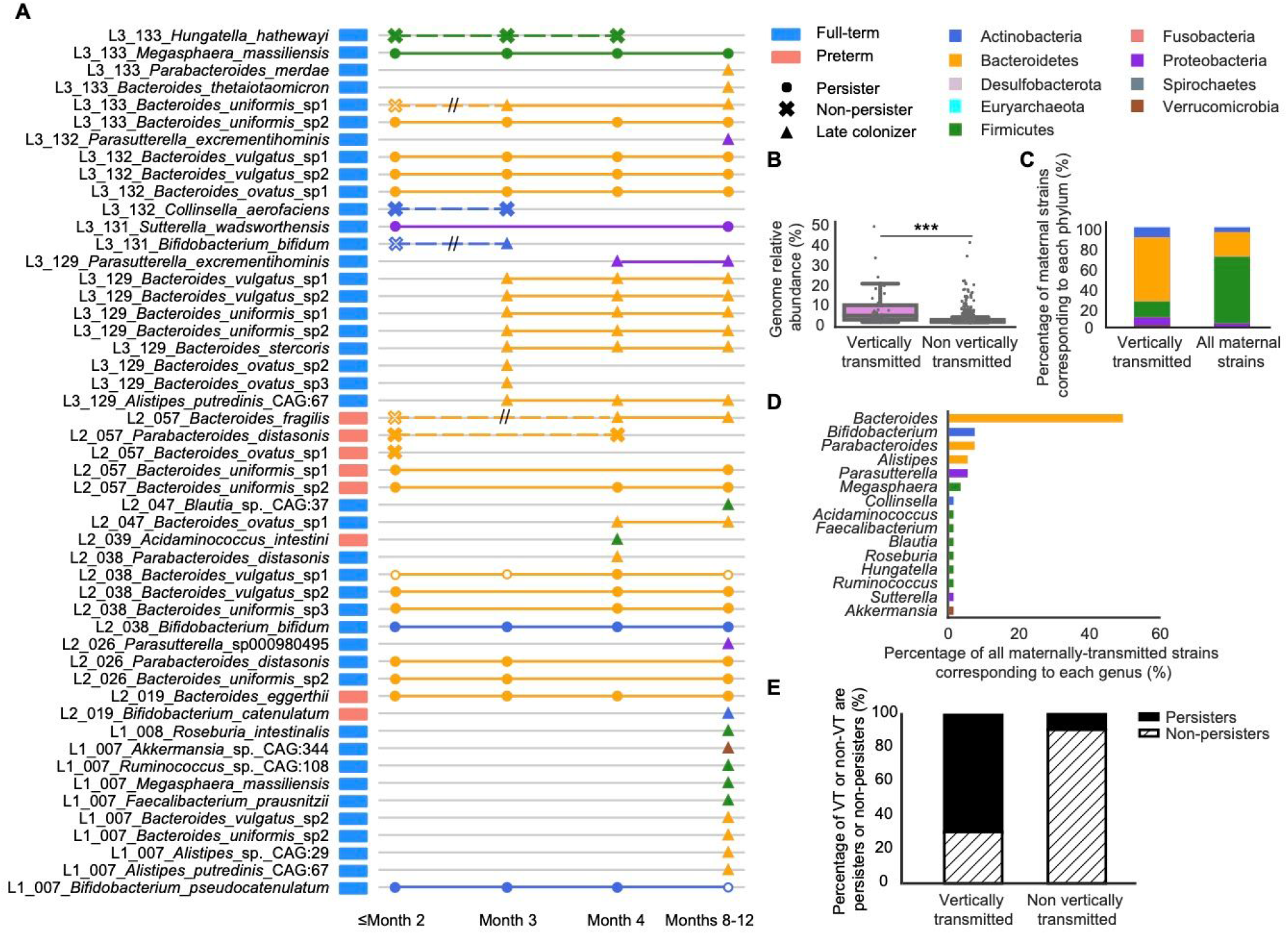
Maternally derived intestinal bacterial strains are more likely to be persisters in the infant gut. (A) Schematic of all 50 strains that were maternally transmitted to infants. Each row represents an infant-specific, maternally transmitted strain, and marks represent months in which the strain was detected. Shapes represent distinct strain identities (i.e., persisters, non-persisters and late colonizers). Solid marks indicate ≥99.999% popANI between the infant and mother strains, and hollow marks indicate windows in which the identity of the infant and mother strains fell below the strain cutoff. Non-persisters of the same subspecies are connected via dashed lines whereas persisters or late colonizers of the same subspecies are connected via solid lines. Double hashing indicates a non-persister early colonizer that is not maternally derived being replaced by a closely related maternal intestinal strain. (B) Relative abundances of maternal subspecies that were and were not vertically transmitted to the infant gut microbiomes. Each dot represents a subspecies detected from a maternal fecal sample. (C) Phylum-level taxonomy of strains that were maternally transmitted to infants as well as all maternal intestinal strains. (D) Percentage of maternally transmitted strains by genus and is colored by phylum. (E) Percentage of persisters (solid black) and non-persisters (dashed) derived from the maternal gut microbiomes and other sources.

The strains that were vertically transmitted were significantly more abundant in the maternal gut microbiomes than strains that were not passed on to infants (*P* = 2.4e-16, Wilcoxon rank-sum test) (Figure 3B). Correspondingly, maternally acquired strains were also more abundant than noninherited strains in the infant gut microbiomes across all time windows (*P* < 0.001, Wilcoxon rank-sum test with FDR correction). Regardless of gestational age or delivery mode, Bacteroidetes were significantly enriched and Firmicutes were significantly depleted (*P* = 2.4e-09 and 2.4e-13, respectively; Fisher’s exact test with FDR correction) among maternally transmitted strains (Figure 3A,C). At the genus level, *Bacteroides* and *Parasutterella* were more likely to be acquired by infants from their mothers than other bacterial genera (*P* = 2.0e-08 and 0.028, respectively; Fisher’s exact test with FDR correction) (Figure 3A,D). *B. uniformis* and *B. vulgatus* were the two most commonly observed species to be maternally transmitted in this cohort (*P* = 3.7e-05 and 0.0015, respectively; Fisher’s exact test with FDR correction). We did not detect significant differences in vertical transmission when segregating infants by various clinical variables (i.e., gestational age, feeding). However, when considering the 12 cases of maternal-infant strain sharing, we found that mothers who delivered infants via cesarean delivery (C-section) transmitted a significantly higher percentage of their gut microbiomes to infants than mothers who delivered vaginally (*P* = 0.014, Wilcoxon rank-sum test).

Maternally transmitted strains were found to be significantly more likely to be persisters in the infant gut microbiomes than strains derived from other sources, suggesting strains acquired from the maternal gut microbiomes are likely to be well adapted to the infant gut (*P* = 4.0e-11, Fisher’s exact test) (Figure 3A,E). These maternally transmitted persisters were primarily Bacteroidetes, whereas persisters not detected in maternal fecal samples were mostly Firmicutes and Actinobacteria. Importantly, we detected new strains being transmitted from mothers to the infant gut microbiomes throughout the first year of life, suggesting vertical transmission is not limited to the intrapartum or postpartum time periods (Figure 3A).

### Non-related infants rarely shared bacterial strains

In addition to examining strain persistence and maternal strain transmission, we also searched for strain sharing between different infants in the study. When considering all possible pairs of individual infants, 18 of 231 infant pairs shared at least one bacterial strain (Figure 4A). While the majority of infant pairs shared no more than two bacterial strains, full-term infants 7 and 133 shared 11 strains (Figure 4). Our decontamination analysis ensured that this was not a result of cross-sample contamination. We therefore hypothesized, and later confirmed by searching medical record data, that these two de-identified infants were siblings, with infant 7 being born two years earlier. We examined the gut microbiomes of the siblings and their mother in greater detail in the next section. No other infants in our study were biologically related.

**Figure 4.**
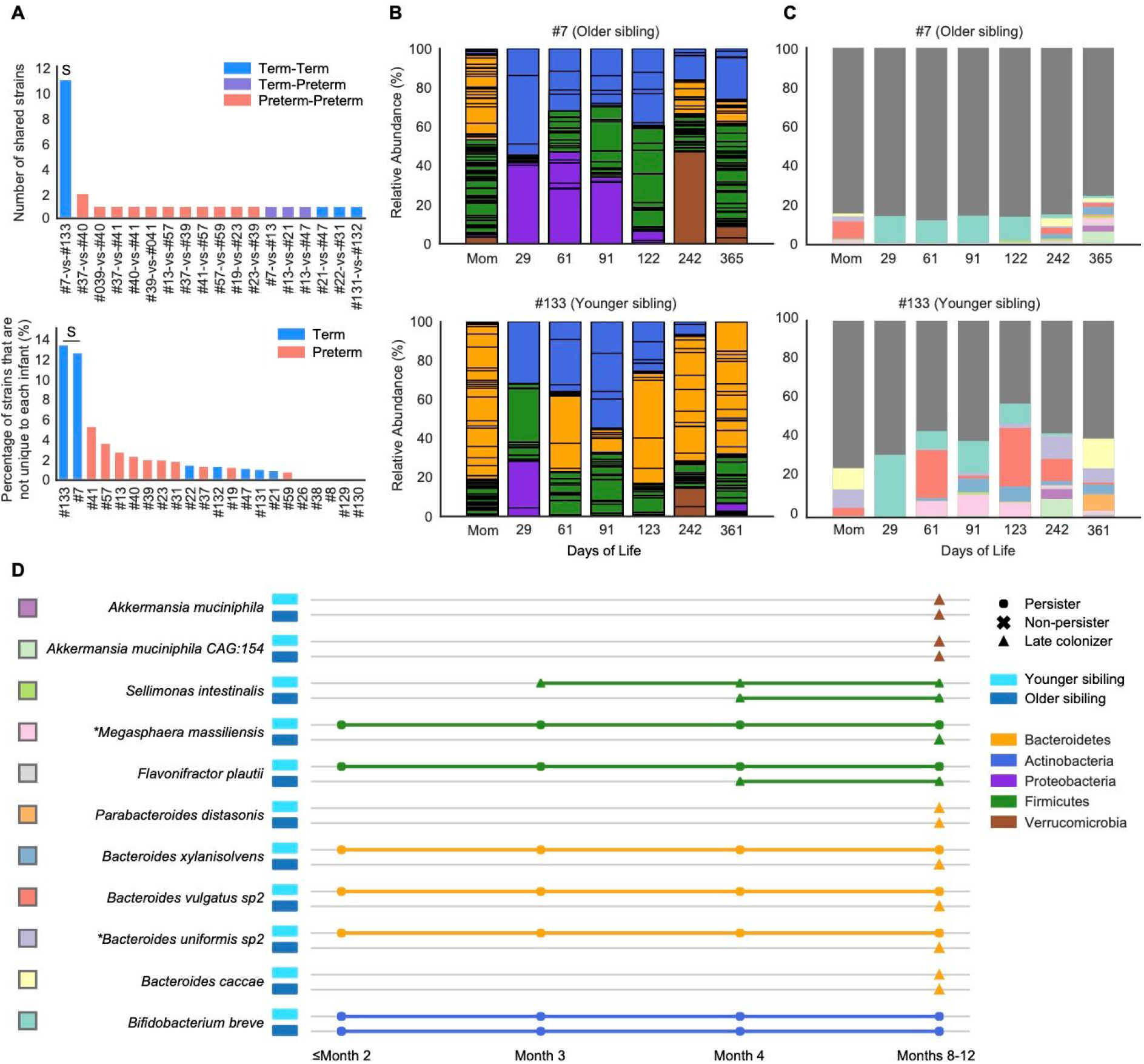
A pair of siblings shared significantly more bacterial strains than non-related infants. (A) (Top) The number of strains that were shared between infant pairs. (Bottom) The percentage of the infant gut microbiome that shared near-identical (≥99.999% popANI) strains with other infants. “S” in both panels refers to the infants 7 and 133 sibling pair. (B) Overview of year-1 gut microbiome compositions of the siblings. Bar height represents normalized subspecies relative abundance, and bars are colored by phylum. Sections of the same color with horizontal black lines correspond to individual subspecies of the same phylum. All maternal fecal samples were grouped into “Mom” on the x-axis. (C) Normalized relative abundance of the 11 strains shared between the siblings throughout the first year of life. Each shared strain is assigned to a unique color and the rest of strains were all colored in dark gray. (D) Longitudinal detection of the 11 strains shared between the siblings. Colored squares on the left of the species names correspond to Figure 4C barplot color scheme. Each row represents a sibling shared strain and is colored based by phylum. Shapes represent distinct strain identities (i.e., persisters, non-persisters and late colonizers). Older and younger siblings are represented by darker and light blue rectangles, respectively (* = maternally transmitted strains).

After excluding the single sibling pair, preterm infants were far more likely to share strains with other preterm infants than full-term infants were to share strains with other full-term infants (*P* = 4.6e-04, Fisher’s exact test) (Figure 4A). Most sharing among preterm infants occurred before the infants were discharged from the hospital, pointing to the hospital environment as a potential strain source. *C. butyricum* was the most widely shared species among preterm infants, and one *C. butyricum* strain was shared by ≥5 non-related preterm infants based on pairwise comparisons.

### A pair of siblings shared a significant number of strains throughout their first year of life

To further investigate strain sharing in the sibling gut microbiomes, we closely examined the gut microbial communities of full-term infants 7 and 133 and the two fecal samples provided two years apart by their mother (Figure 4B-D). The two siblings were both born via C-section and were breastfed exclusively before weaning. During the first year of life, when compared to the older sibling, the younger one had fewer Proteobacteria and Verrucomicrobia subspecies and more Bacteroidetes. We speculated that changes in the gut microbiome of the mother around the time of birth of the second compared to the first child might explain the observed compositional differences between the siblings. Indeed, the mother’s gut microbiome was nearly twice as enriched in Bacteroidetes and contained about six times less abundance of Proteobacteria and no Verrucomicrobia around the time of second delivery compared to the first (Figure 4B).

The 11 bacterial strains that were shared by the siblings accounted for ~20% of the overall gut microbiome of the older sibling and ~50% of the gut microbial community of the younger sibling (Figure 4C). Interestingly, only one of the 11 shared strains (*B. breve*), not maternally derived, persisted in both siblings throughout most of their first year of life, and five of the 11 shared strains were classified as persisters only in the younger sibling (Figure 4D). No other infant pairs shared any bacterial persisters. Since most shared strains were late colonists in the older sibling but were early colonists in the younger sibling and they were mostly not detected in the gut microbiome of their mother, we hypothesize that strains may have been transmitted from the older to the younger sibling (Figure 4D).

Having collected two fecal samples from the same mother also allowed us to search for bacterial strains present in both samples. Of the 99 and 94 subspecies detected from the first and second maternal fecal samples, respectively, 12 (mostly Bacteroidetes) shared ≥99.999% popANI. Notably, these 12 strains constituted 20% of the maternal gut microbiome at the time of birth of her first child and ~50% of her gut microbiome two years later (Figure S2A). Of these 12 strains, a *B. uniformis* and a *Megasphaera massiliensis* strain were acquired by both siblings. Both of these strains were persisters in the younger sibling. Two of the other 10 maternal strains were detected when the younger sibling was one-year-old; another one of the 10 strains was detected in the older sibling at age one (Figure S2B).

### Diverse carbohydrate active enzymes are implicated in *Bifidobacterium* and *Escherichia* persistence

We next investigated whether specific capacities of early colonizers are associated with strain persistence. To do this, we compared the gene content of persisters and non-persisters during the critical first two months of life where strain outcomes were determined (Methods; Tables S3,4).

As the ability to metabolize a variety of carbohydrates is considered to be important for surviving in the gut (El Kaoutari et al., 2013; Fischbach and Sonnenburg, 2011), we hypothesized that persister genomes would be enriched with carbohydrate active enzymes (CAZymes) when compared to non-persisters. To test our hypothesis, we annotated genes encoding CAZymes (Methods) and measured the diversity of glycoside hydrolases (GHs), polysaccharide lyases (PLs), carbohydrate esterases (CE), and carbohydrate-binding modules (CBMs) in the genomes of persisters and non-persisters. GHs, PLs, CEs and CBMs were chosen because they are directly involved in carbohydrate degradation (El Kaoutari et al., 2013).

Overall, persisters had a significantly higher CAZyme Shannon diversity than non-persisters (*P* = 1.1e-08; Wilcoxon rank sum test). However, persisters in our study were primarily *Bacteroides* and *Bifidobacterium*, whose genomes are known to densely encode glycan degradation genes (Marcobal et al., 2011; Sela and Mills, 2010; Wexler and Goodman, 2017). To address this potential taxonomy bias, we restricted our comparisons of CAZyme diversity to be between persisters and non-persisters from the same genus or species. Further, we required at least three persister and three non-persister strains for comparisons to retain statistical power. Of the five genera (*Escherichia*, *Bifidobacterium*, *Klebsiella*, *Streptococcus* and *Bacteroides*) meeting these criteria, *Escherichia* and *Bifidobacterium* persisters encoded a higher CAZyme Shannon diversity when compared to their corresponding non-persisters (*P* = 0.0060 and 0.025, respectively; Wilcoxon rank-sum test). *E. coli*, which had 3 persisters and 25 non-persisters, was the only species that passed the filtering criteria, and its persisters encoded a significantly higher Shannon diversity of CAZymes than non-persisters (*P* = 0.0060; Wilcoxon rank-sum test).

We next searched for specific CAZymes that were enriched in *Bifidobacterium* and *Escherichia* persisters (Methods). Of the 96 CAZymes examined in *Bifidobacterium* persisters, three (GH1, CBM41, and CE4) were significantly enriched among persisters (*P* < 0.05, Fisher’s exact test with FDR correction), and they were all predicted to participate in metabolizing dietary polysaccharides (Table S3). In *E. coli*, 7 out of 43 CAZymes examined were significantly enriched in persisters (*P* < 0.05, Fisher’s exact test with FDR correction). Most CAZymes such as GH33, PL9_1 and GH65 that were enriched in *E. coli* persisters involved in activities such as metabolizing small molecules including sugar byproducts of mucin and dietary polysaccharides degradation carried out by other community members (Table S3). Interestingly, GH153, a CAZyme predicted to be involved in biofilm formation (Little et al., 2018), is also enriched in *E. coli* persisters, suggesting *E. coli* persisters might carry other traits, in addition to carbohydrate metabolism, that enable its stable colonization in the gut.

### Surface attachment and iron acquisition contribute to *E. coli* persistence

To identify other functions besides carbohydrate metabolism that could contribute to *E. coli* persistence in the infant gut, we compared the gene content of *E. coli* persisters and non-persisters present during the first two months of life using the Kyoto Encyclopedia of Genes and Genomes (KEGG), Pfam, and Transporter Classification (TC) databases (Methods).

Of the KEGG Orthologies (KOs), Pfams and TC numbers (TC#) examined, 64 KOs, 100 Pfams and 24 TC# were significantly enriched in *E. coli* persisters (*P* < 0.05; Fisher’s exact test with FDR correction). These were associated with 267 genes in total. No annotations were significantly enriched in non-persisters. Notably, 4 KOs, 18 Pfams and 4 TC# were present in all *E. coli* persisters and completely absent in all *E. coli* non-persisters. These were all linked to four genes (Tables 1 and S4). Two of these were CdiB/CdiA proteins and antigen 43, both of which have been reported to be involved in surface adhesion (i.e., via cell-cell aggregation and/or enhancing biofilm formation) (Ruhe et al., 2015; Trunk et al., 2018). The other two were GatA and thioesterase, both of which are predicted to be part of the bacterial toxin colibactin biosynthesis gene cluster. Colibactin damages intestinal epithelial cells, and could increase intestinal permeability to enhance the translocation of *E. coli* strains (Secher et al., 2015).

**Table 1.** Selected annotations that were enriched in *E. coli* persisters

| Annotation definition | KO(s) | Pfam(s) |
| --- | --- | --- |
| Surface adhesion |  |  |
| Antigen 43 | K12687* |  |
| Serine protease autotransporter | K12648* | PF02395* |
| CdiB/CdiA proteins (Putative filamentous hemagglutinin) | K15125* | PF05860*,PF13332*,<br>PF04829*,PF15530*,<br>PF03865*,PF08479*,<br>PF17287* |
| Type VI secretion system | K11891*,K11892*,<br>K11893*,K11895*,<br>K11896*,K11900*,<br>K11901*,K11903*,<br>K11904,K11905*,<br>K11906*,K11907*,<br>K11910*,K11911* | PF05488*,PF05591*,<br>PF05936*,PF05943*,<br>PF05947*,PF06744*,<br>PF06761*,PF06812*,<br>PF06996*,PF09850*,<br>PF10106*,PF12486*,<br>PF12790*,PF13296*,<br>PF16989* |
| Biofilm synthesis | K11931*,K11935*,<br>K11936*,K11937* | PF13994*,PF14883* |
| Iron acquisition |  |  |
| Yersiniabactin biosynthesis | K04781,K04783,<br>K04784*,K04785,<br>K04786,K05372*,<br>K05373,K05374*,<br>K15721 | PF08242,PF08659,<br>PF16197 |
| Aerobactin biosynthesis | K03894*,K03895*, | PF04183*,PF13434*, |
|  | K03896*,K03897* | PF13523* |
| Manganese/iron transport system, SitABCD | K11604*,K11605*,<br>K11606*,K11607* |  |
| Bacterial toxins |  |  |
| Uropathogenic Escherichia coli Colicin-Like (Usp) |  | PF01320*,PF05638*<br>PF06958* |
| Colibactin biosynthesis | K01071*,K01426* | PF01527*,PF08659,<br>PF13602*,PF13683*,<br>PF14765*,PF16197,<br>PF01425*,PF08020*,<br>PF16197 |

\* = q-value < 0.05; annotations without “\*” are those that had *P*-values <0.05 prior to correcting for multiple comparisons but not after.

We also identified genes for functions that were enriched, but not exclusively present in persisters. Many of these are also involved in surface adhesion (e.g., type VI secretion system and biofilm biosynthesis). Also enriched was uropathogenic *Escherichia coli* colicin-like protein (Usp) that has been shown to cause damage to the eukaryotic cells (Nipič et al., 2013). Other enriched traits include iron acquisition (e.g., siderophore production and manganese/iron transporters) and sugar and amino acid metabolism (e.g., pectin-associated metabolism and D-serine detoxification and metabolism) (Tables 1 and S4). *E. coli* persisters dedicated significantly higher percentages of their genomes to surface adhesion and iron acquisition (*P* = 0.0067 and 0.017, respectively; Wilcoxon rank-sum test) than non-persisters, suggesting that these traits are particularly important for persistence of *E. coli* early colonizers in the infant gut.

We used comparative genomic analyses to verify that genes that are apparently absent or relatively uncommon in *E. coli* non-persisters were not simply missed due to missing genome fragments (Methods). As expected, we detected insertions/deletions involving enriched/absent genes in otherwise syntenous regions. For instance, we found that genes involved in synthesis of the colicin-like protein (Usp) and its associated immunity protein, as well as a large region that encodes a Type VI secretion system, were absent in otherwise syntenous regions of the persister and non-persister *E. coli* genomes (Figure S3).

### Divergent early gut microbiomes of full-term and preterm infants converged by age one

To understand how early-life gut microbiome assembly might differ between full-term and preterm infants at the community-level, we measured the alpha- and beta-diversity of the two infant groups using the Shannon index and the UniFrac distance (Lozupone and Knight, 2005), respectively (Methods).

Shannon indices increased over time across all infants (Figure S4A). However, the gut microbiome of full-term infants had a higher Shannon diversity during the first three months of life (*P* = 0.037, Wilcoxon rank-sum test) than preterm infants. During this time period, preterm infants’ gut microbiomes were disproportionately dominated by bacteria that are common in the hospital environment, including members of ESKAPE pathogens such as *Klebsiella pneumoniae*, *Enterococcus faecium* and *Enterobacter* spp. (Rice, 2008) (Figure 5A). Full-term infant microbiomes over the first three months were more taxonomically balanced than those of preterm infants. Although some ESKAPE organisms were also present in full-term infants, their gut microbiomes contained more organisms that normally considered to be commensals, such as *Bacteroides* and *Bifidobacterium*. From month 4 onwards, the alpha-diversity differences between the gut microbiomes of full-term and preterm infants decreased. In preterm infants, a reduction in the relative abundances of Proteobacteria subspecies was coupled with increases in abundance of Actinobacteria and Bacteroidetes subspecies (Figure 5A). By age one, both full-term and preterm infants’ gut microbiomes were dominated by Firmicutes and Bacteroidetes. However, the gut microbiomes of full-term infants had higher percentages of Actinobacteria than those of preterm infants (Figure 5A).

**Figure 5.**
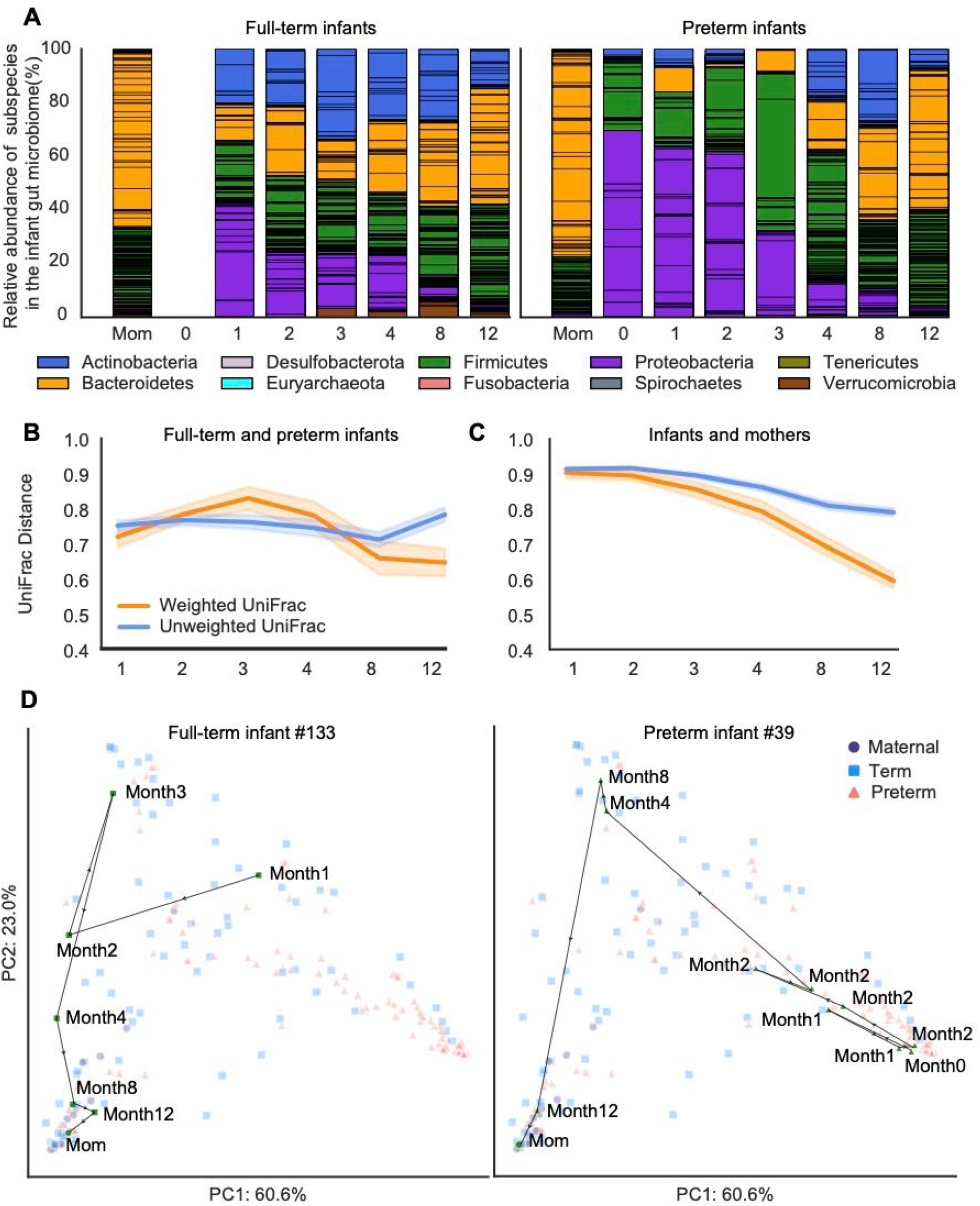
Community assembly dynamics of full-term and preterm infant gut microbiome during the first year of life. (A) Overview of year-1 gut microbiome compositions of the full-term and preterm infants. Bar height represents normalized subspecies relative abundance, and bars are colored by phylum. Sections of the same color with horizontal black lines correspond to individual subspecies of the same phylum. X-axis represents months of life and all maternal fecal samples were grouped into “Mom” on the x-axis. (B-C) Weighted (orange) and unweighted (blue) UniFrac comparing the compositional changes of the gut microbiomes of full-term and preterm infants (B) as well as of infants and mothers (C). X-axis represents months of life. (D) Gut microbiome assembly trajectories of a representative full-term (left) and a representative preterm (right) infant. Shapes and colors represent distinct sample types (preterm, full-term or maternal fecal samples). Samples belonging to the representative infant are colored green in order to distinguish them from samples from other infants, which are colored based on sample types. PCA is based on weighted UniFrac distance.

Weighted and unweighted UniFrac distances were used to measure beta-diversity in gut microbiomes of term and preterm infants. Weighted UniFrac, which considers the relative abundances of individual taxa, indicated that the gut microbiomes of preterm infants diverged from those of full term infants between months 1 and 3. However convergence between the gut microbiomes of preterm and full-term infants began at month 3 and accelerated between months 4 and 8 (*P* = 2.0e-04, Wilcoxon rank-sum test). Overall, weighted UniFrac inidcated that the microbiomes of full-term and preterm infants converged by age one (*P* = 0.0024, Wilcoxon rank-sum test) (Figures 5B and S4B). Unweighted UniFrac, which excludes relative abundance, indicated that the gut microbiomes of preterm and full-term infants became similar between months 1 and 8 (*P* = 0.0059, Wilcoxon rank-sum test) but diverged rapidly after month 8. Altogether, unweighted UniFrac suggested that the preterm and full-term infant microbiomes became more distinct by age one (*P* = 0.014, Wilcoxon rank-sum test) (Figures 5Band S4B). To evaluate the contrasting outcomes of the two UniFrac distances, we also tested for convergence of gut microbiomes of full-term and preterm infants using Bray-Curtis distance (Method) (Figure S4C). Consistent with weighted UniFrac, Bray-Curtis distance indicated that the gut microbial compositions of full-term and preterm infants largely converged by age one (*P* = 1.7e-31, Wilcoxon rank-sum test).

To further examine the maturation of the infant gut microbiome, we measured the beta-diversity between the gut microbiomes of infants and mothers. Both weighted and unweighted UniFrac distances showed a gradual convergence between infants and mothers. However, unweighted UniFrac indicated a slower convergence rate than weighted UniFrac (Figures 5C and S4B).

Development of gut microbiomes of preterm and full-term infants in the context of maternal microbiome composition was further examined via principal component analysis (PCA) (Methods). Each infant’s assembly trajectory was visualized in a PCA by tracking the changes in composition between consecutive sampling time points (Figure 5D). The gut microbiomes of preterm infants, full-term infants and their mothers formed distinct clusters in PCA space (permutational multivariate analysis of variance (PERMANOVA), *P* < 0.001, 1000 permutations, weighted UniFrac distance) (Figure S4D). However, over time, the infant gut microbiomes all moved toward the PCA region where the maternal samples were placed. Indeed, chronological age played a significant role in driving the gut microbiome changes for both full-term and preterm infants (PERMANOVA, *P* = 0.040 and *P* < 0.001, respectively, 1000 permutations, weighted UniFrac distance). Notably, the trajectories of preterm infant microbiomes were significantly different from those for full-term infants, possibly because their initial gut microbiomes were more distinct from the maternal gut microbiomes than those of full-term infants. In fact, Jaccard dissimilarity comparing consecutive fecal metagenomes of each infant (Methods) indicated that the changes between the early and late gut microbiomes of preterm infants were significantly larger than full-term infants (*P* = 0.0092; Wilcoxon rank-sum test).

## DISCUSSION

We conducted strain-resolved metagenomics analyses to investigate ecological succession in the early-life gut microbiome. We showed that gut microbiome assembly is a largely individualized process as infants rarely share identical strains over a long period of time. Our finding that 11% of bacterial colonizers persist through the first year of life extends results of prior studies that used primarily isolation based strategies to show that specific bacterial strains can persist in gut microbiomes for years (Faith et al., 2013; Nowrouzian and Oswald, 2012; Zhao et al., 2019). Our use of genomes derived from the metagenomes of the study cohort enabled detection of the actual strains present and ensured more complete gene inventories and robust SNP detection. Further, our exceptionally stringent strain definition method provided high confidence that strains classified as persisters were not closely related strains that were introduced into the gut microbiome at a later time point.

Our results showed that the majority of the initial gut microbiome is transient, and only a small percentage of the early colonists persist until age one. High strain turnover is not surprising because processes such as maturation of the immune system and events including dietary shifts (i.e., the cessation of breast milk) should impose changing selective pressures on members of the infant gut microbiome (Bäckhed et al., 2015). The persistence of some early arrived strains may in part be due to a priority effect in which they shape the trajectory of the developing infant microbiome (e.g., by taking space and/or resources that would otherwise be available to later arriving strains or by controlling the local environment).

We identified one important factor that seems to dictate whether a colonizing strain is in the small subset of strains that persist over the first year of life. By analyzing maternal fecal samples collected around the time of birth, we determined that strains derived from the maternal gut such as *Bacteroides* are significantly more likely to colonize the infant gut long-term than non-inherited strains. This finding extends previous work conducted over a shorter period of time, using less stringent SNP detection and mapping to clade-specific genes (Ferretti et al., 2018), and a study that relied on analysis of rare SNPs of abundant species only (Korpela et al., 2018). In combination, these results could indicate that maternally acquired intestinal strains are better adapted to the gut than strains from other sources.

We determined that specific bacterial taxa are far more likely than others to persist in the developing infant gut. We speculate that early-life diet, especially breast milk, favors the growth of certain commensal bacteria such as *Bacteroides* and *Bifidobacterium* (Marcobal et al., 2011; Sela and Mills, 2010). Indeed, we found that *Bacteroides* and *Bifidobacterium* strains, many of which can metabolize human milk oligosaccharides (HMOs) from breast milk, are most likely to persist in the infant gut microbiome and we also show that infants who were breastfed only had more bacterial persisters than those who had a mixture of breast milk and formula. This finding is in line with the hypothesis that breast milk, which includes components such as HMOs and immunoglobulins that select for certain bacteria over others, offers the evolutionary benefit of promoting the stable colonization of beneficial bacteria in the infant gut (Doare et al., 2018; Wang et al., 2020).

The ability to metabolize HMOs alone may not be sufficient to confer strains with the ability to persist in the infant gut, since all infants in this study were weaned from maternal milk or formula before age one. Indeed, we found that *Bifidobacterium* persisters encode a significantly higher diversity of CAZymes than *Bifidobacterium* strains that failed to persist during the first year of life. Notably, CAZymes that are enriched among *Bifidobacterium* persisters are mostly involved in degrading dietary polysaccharides such as plant cell wall materials, a finding consistent with previous work (Bäckhed et al., 2015; Koenig et al., 2011). The enrichment of CAZymes dedicated to degrading dietary carbohydrates suggest the metabolic flexibility of *Bifidobacterium* persisters, enabling adaptation as the infant’s diet changes. This is consistent with prior work that proposed a link between plant polysaccharide metabolic capacity and the ability of a strain to adapt by shifting metabolism following introduction of solid food (Fischbach and Sonnenburg, 2011). We were not able to detect differences in carbohydrate metabolism between *Bacteroides* persisters and non-persisters, although differences may be detected in future studies that include larger datasets.

A diverse pool of CAZymes was also identified in *E. coli* persisters, which were only found in preterm infants in our study. However, investigation of genes enriched in *E. coli* persisters revealed additional traits that may offer fitness advantages, many of which are common to *E. coli* pathogens including uropathogenic *E. coli* (UPEC). Specifically, the genes common to pathogens and also present in majority of persisters include antigen 43 (Trunk et al., 2018; Ulett et al., 2007; van der Woude and Henderson, 2008), serine protease autotransporters (Habouria et al., 2019; Ruiz-Perez and Nataro, 2014), siderophore biosynthesis (Garcia et al., 2011; Schubert et al., 2002; Watts et al., 2012) and CdiB/CdiA proteins (Aoki et al., 2005; Ruhe et al., 2015). In the cohort studied here, one *E. coli* persister was detected in a preterm infant who survived two necrotizing enterocolitis (NEC) events. This observation may relate to findings of Ward et al., who reported that intestinal colonization by UPEC strains is a significant risk factor for the onset of NEC in infants (Ward et al., 2016). Even more suggestive of a role in disease, persister *E. coli* strains were detected in fecal samples of two other infants who had late onset sepsis (LOS) prior to the onset of disease.

A recent comparison of the gut microbiomes of preterm and near-term infants over 21 months via mapping to clade-specific marker genes found evidence for convergence, but noted persistence of antibiotic resistance genes in preterm infants (Gasparrini et al., 2019). The authors also isolated *Enterobacteriaceae* to show persistence of some genotypes. Using cultivation-independent genome-resolved strain analyses, we also found that initially distinct gut microbiomes of preterm and full-term infants largely converged by the time the infants were one-year-old. However, strains of *E. coli* carrying virulence genes persisted only in preterm infants. In combination, these studies imply that prematurity can impact the infant microbiome for a span of time sufficient to have an impact on infant development.

It is critical to understand the factors that lead to persistence of early acquired bacterial strains, especially for preterm infants, as these strains have the potential to impact infant health and development over the first year of life, and potentially beyond. Of specific concern are strains that are predicted to be pathogenic, such as the *E. coli* persisters we detected in preterm infants. Genome resolution allowed us to show that strain source, strain phylogeny and genetic traits that enable long term gut microbiome residency. Taken together, our findings provide clues to the development of microbiome therapies.

## ACKNOWLEDGMENTS

We thank Rohan Sachdeva, Ka Ki Lily Law and Shufei Lei for the technical support, Raphaël Méheust and Jacob West-Roberts for assistance in bioinformatics tools, Alexandra Sheppeck for fecal sample collection and Adair Borges for comments on the manuscript. We are also grateful for all the families that participated in this study. For funding support, we acknowledge NIH award RAI092531A to JFB and MJM and Chan Zuckerberg Biohub support to JFB.

## AUTHOR CONTRIBUTIONS

Y.C.L., M.R.O., M.J.M., and J.F.B. designed the study; B.A.F. performed DNA extractions from all fecal samples; R.B. supervised the enrollment of infants; Y.C.L. coordinated the acquisition of the metagenomics data; Y.C.L and J.F.B. wrote the manuscript and all authors contributed to the manuscript revisions.

## METHODS

### Details of infant recruitment

This study was reviewed and approved by the University of Pittsburgh Institutional Review (IRB STUDY19120040). To investigate the gut microbiomes of premature and full term infants over the first year of life, we recruited a cohort of 183 infants. However, withdrawal from our study, missing samples at key timepoints or low sample biomass led to exclusion of many infants from this study. Ultimately, we acquired longitudinal samples from 23 full-term and 19 preterm from birth to age one (Figure S1). Fecal samples from enrolled infants and their mothers were all collected at the UPMC Magee-Womens Hospital (Pittsburgh, PA) over the course of three years. While full-term infants were discharged from the hospital within 3 days after birth and received no perinatal antibiotics, all preterm infants received empiric antibiotics immediately following birth during an evaluation for early-onset sepsis and then spent their first 2-3 months in the hospital. In addition to infant fecal samples, we collected a single fecal sample from 28 mothers of 29 infants within the first two weeks after delivery. All samples were collected with parental consent and subjects were de-identified before the receipt of samples. Genome-resolved metagenomics analyses in this study were performed on 13 out of 23 full-term and 9 out of 19 preterm infants due to well-to-well contamination on samples from 20 out of 42 infants. De-identified metadata for the 22 infants whose samples were not contaminated is provided in Tables S2.

### Sample collection and metagenomic sequencing

Throughout the first year of life, infant fecal samples were collected either at UPMC Magee-Womens Hospital by trained nurses or at home by parents provided with detailed collection instructions. Specifically, fresh infant stool samples were collected directly from infants while they were actively excreting or from diapers shortly after the stools were released. Maternal fecal samples were collected using a commode specimen collector, from which fecal samples were transferred into a collection tube. All stool samples collected at the hospital were immediately stored at −80°C following collections. Samples collected at home were stored in home freezers until they were picked up by research staff and transferred to the −80°C condition. DNA extraction of frozen fecal samples was performed via the Qiagen DNeasy PowerSoil HTP 96 DNA isolation kit with modifications to the manufacturer’s protocol. For each 96-well extraction plate, a reagent-only negative control was included.

Metagenomic sequencing of collected infant and maternal fecal samples was performed in collaboration with the California Institute for Quantitative Biosciences at UC Berkeley (QB3-Berkeley). Library preparation on all samples was performed as previously described (Olm et al., 2019b). Final sequence ready libraries were pooled into 2 subpools and visualized and quantified on the Advanced Analytical Fragment Analyzer. Four samples did not fit nicely into either subpool so their libraries were quantified separately. All libraries were then evenly pooled into a single pool and checked for pooling accuracy by sequencing on Illumina MiSeq Nano sequencing runs. The single pool was adjusted based on MiSeq sequencing run and sequenced on individual Illumina NovaSeq6000 150 paired-end sequencing lanes with 2% PhiX v3 spike-in controls. Post-sequencing bcl files were converted to demultiplexed fastq files per the original sample count with Illumina’s bcl2fastq v2.20 software.

### Metagenomic assembly and gene prediction

Reads from all 402 samples were trimmed using Sickle (www.github.com/najoshi/sickle), and reads that mapped to the human genome with Bowtie2 (Langmead and Salzberg, 2012) under default settings were discarded. Reads from each sample were then assembled independently using IDBA-UD (Peng et al., 2012) under default settings. Co-assemblies were also performed for each infant, in which reads from all samples of that infant were combined and assembled together. Scaffolds that are <1 kb in length were discarded. On average, 93.2% of the sequencing reads (95% confidence interval, 92.4%-94.4%) were *de novo* assembled into scaffolds ≥1 kbp in length per sample. Remaining scaffolds were annotated using Prodigal (Hyatt et al., 2010) to predict open reading frames using default metagenomic settings.

### Metagenomic *de novo* binning

Pairwise cross-mapping was performed between all samples from each infant to generate differential abundance signals for binning. Each sample was binned independently using three automatic binning programs: metabat2 (Kang et al., 2019), concoct (Alneberg et al., 2014) and maxbin2 (Wu et al., 2016). DasTool (Sieber et al., 2018) was then used to select the best bacterial bins from the combination of these three automatic binning programs. The resulting draft genome bins were dereplicated at 98% whole-genome average nucleotide identity (gANI) via dRep (v2.6.2) (Olm et al., 2017b), using a minimum completeness of 75%, maximum contamination of 10%, the ANImf algorithm, 98% secondary clustering threshold, and 25% minimum coverage overlap. Genomes with gANI ≥98% were classified as the same subspecies, and the genome with the highest score (as determined by dRep) was chosen as the representative genome from each subspecies. A total of 1005 genomes were selected to represent unique microbial “subspecies” and they had an average of 96% completeness and 1.05% contamination.

### Taxonomy assignment

The amino acid sequences of predicted genes of all assembled bins were searched against the UniProt100 database using the usearch ublast command with a maximum e-value of 0.0001. tRep (https://github.com/MrOlm/tRep/tree/master/bin) was used to convert identified taxIDs into taxonomic levels. Briefly, for each taxonomic level (species, genus, phylum, etc.), a taxonomic label was assigned to a bin if ≥50% of proteins had best hits to the same taxonomic label. GTDB-Tk was used to resolve taxonomic levels that could not be assigned by tRep (Chaumeil et al., 2019).

### Detection of subspecies and identification of strains using inStrain

Reads from each individual fecal sample were mapped to all 1005 representative subspecies (generated via dRep as described above) concatenated together using Bowtie2 under default settings. inStrain (v1.3.4) *profile* (Olm et al., 2021) was run on all resulting mapping files using a minimum mapQ score of 0 and insert size of 160. Genomes with ≥0.5 breadth (meaning at least half of the nucleotides of the genome are covered by ≥1 read) in samples were considered to be present. inStrain *compare* was used under default settings to compare read mappings to the same genome in different pairs of samples. Samples were considered to share the same strain of the examined genome if the compared region of the genome from samples shared ≥99.999% population-level ANI (popANI). Only genomic areas with at least 5x coverage in samples were compared, and sample pairs with less than 50% of comparable regions of the genome were excluded (≥0.5 percent_genome_compared).

### Genome metabolic annotation

Kyoto Encyclopedia of Genes and Genomes (KEGG) orthology groups (KOs) were assigned to predicted ORFs for all fecal metagenomes using KofamKOALA (Aramaki et al., 2019). Carbohydrate active enzymes (CAZymes) were assigned to all nucleotide sequences using run_dbcan.py (https://github.com/linnabrown/run_dbcan) against the dbCAN HMM (v9), DIAMOND (v0.9.31), and Hotpep (v2.0.8) databases with default settings. Final CAZyme domain annotations were the best hits based on the outputs of all three databases. Domains were also predicted using hmmsearch (v.3.3) (e-value cut-off 1 × 10^−6^) against the Pfam r32 database (El-Gebali et al., 2019). The domain architecture of each protein sequence was resolved using cath-resolve-hits with default settings (Lewis et al., 2019). The transporters were predicted both hmmsearch (same settings as the pfam prediction and domain architecture was resolved using cath-resolve-hits) and BLASTP (v2.10.0) (keeping the best hit, e-value cutoff 1e-20) against the Transporter Classification Database (TCDB) (downloaded in November 2020) (Saier et al., 2006). SignalP (v.5.0b) (parameters, −f short gram+) was used to predict proteins’ putative cellular localization (Armenteros et al., 2019). Transmembrane helices in proteins were predicted via TMHMM (v.2.0) with default settings (Krogh et al., 2001). Secondary metabolites were characterized using antiSMASH (v5.1.2) with default settings (Blin et al., 2019).

### Identification of sources of contamination

One negative reagent control (NC) was included in each 96-well DNA extraction plate, in which no material was added during the DNA extraction step. In total this study involved five extraction plates labeled P1 to P5. NCs were labeled by the plate number (i.e., NC1 refers to the negative control sample on the extraction plate 1). All five NC samples were subjected to the DNA extraction and sequencing the same as the fecal samples. Subspecies present in NC samples were detected via mapping reads from NC samples to all 1005 representative subspecies as described above. Subspecies detection limit was the same as described above. We found two NC samples (NC3 and NC4) had over 50% of their reads mapped to ~60 out of 1005 representative subspecies. To search for subspecies that was unique to NC samples, we recovered draft genomes from all five NC samples and dRep (settings were the same as described above) was run on these genomes together with the 1,0005 dereplicated genomes recovered from fecal samples. Through this approach, we did not find any subspecies that were unique to NC samples.

Detection of bacterial genomes in the NC3 and NC4 could be a result of index hopping, barcode bleeding, reagent contamination, and/or sample spillover (or “well-to-well contamination”). Since all samples were given Unique Dual Indexes, the observed contamination in NC3 and NC4 were unlikely to be a result of index hopping. We also eliminated the possibility of barcode bleeding by resequencing NC3 and NC4 alone. The possibility for reagent contamination to occur in our case was also unlikely since not only did we fail to detect any bacterial genomes in the rest of three NC samples, but we also did find bacterial strains being shared over 50% of the samples either on the same extraction plates or across all five plates. We therefore hypothesized that the detection of intestinal bacterial genomes in NC3 and NC4 was a result of sample spillover within plates 3 and 4. Using the strain-resolved methods detailed above, we detected strain sharing across the extraction plates 3 and 4, but not with the rest of four plates. Given the reliance of our study on robust and accurate detection of strain sharing, we excluded from analysis all samples from plates 3 and 4.

### Detection of mother-to-infant vertical transmission

For each mother-infant pair, every fecal sample from the infant was compared to its maternal fecal sample to search for identical bacterial strains (≥99.999% popANI and ≥0.5 percent_genome_compared) via inStrain *compare* (described above). A strain was considered to be vertically transmitted if it was shared between the maternal fecal sample and at least one infant fecal sample.

### Persister and non-persister detection

“Beginning-end” and “pairwise” approaches were used to identify persister and non-persister strains among early colonizers. The “beginning-end” approach searched for strains which shared ≥99.999% popANI between the first two months of life (≤ month 2) and the last two sampling windows (around months 8 and 12). 54 persisters and 506 non-persisters were detected using this approach. The “pairwise approach” identified strains which shared ≥99.999% popANI across consecutive month windows (≤ month 2 & month 3, month 3 & month 4, month 4 & ≥ month 8), yielding 36 persisters and 525 non-persisters. These two approaches combined resulted in the total identification of 59 persisters and 501 non-persisters across 22 infants.

We chose to classify strains as persisters using the month 8 cutoff as we did not want to exclude persisters that would be missed due to lack of a month 12 sample (one infant) or poor genome recovery from month 12 samples (eight infants). We chose the cutoff of 99.999% popANI for persistence because we calculated that it is unlikely for a strain to acquire 40 SNPs in one year, given an average bacterial genome size of ~4 Mbp and the expected rate of *in situ* bacterial evolution in the human gut (~0.9 single nucleotide polymorphisms (SNPs)/genome/year (Zhao et al., 2019)).

### Persister and non-persister functional enrichment analysis

Genes from persisters and non-persisters of each examined bacterial group (i.e., *Bifidobacterium* and *E. coli*) were profiled via inStrain *profile* under default settings. Genes were considered to be present if they had ≥1x coverage across ≥50% of their length. Genes were annotated using the CAZy, KEGG, Pfam and Transporter Classification (TC) databases as described above. Fisher’s exact test (as implemented using the Scipy module “scipy.stats.fisher_exact”) followed by false discovery rate (FDR) correction were run on genes annotated with each database (CAZy, KEGG, Pfam and TCDB) independently to identify annotations from each database that were significantly enriched in persisters or non-persisters (*P* < 0.05 with FDR correction). Annotations were further verified using the UniProt100 and UniRef databases. Annotations that were present in more than 50% of all persisters and non-persisters as well as those that were present in less than 50% of all persisters and non-persisters were excluded from this analysis.

To search for other traits besides carbohydrate metabolism that were associated with *E. coli* persistence in the infant gut, genes with annotations that were significantly enriched in *E. coli* persisters or non-persisters from KEGG, Pfam and TC databases were combined. Further, we located genomic positions of these differentially enriched annotations and inferred their genome-specific functions by examining up- and downstream neighboring gene annotations. The final datasheet listing annotations that were differentially enriched in *E. coli* persisters and non-persister is provided in Table S4. Annotations with *P*-values < 0.05 only (q-values > 0.05) are also provided in Table S4.

### Gene-based nucleotide diversity analysis on annotations that were significantly enriched in *E. coli* persisters

Gene-specific nucleotide diversity and coverage were obtained from inStrain *profile* as described above. For each gene with an annotation (KO, Pfam and/or TC#) shown to be enriched in *E. coli* persisters (methods detailed above), we compared its nucleotide diversity on the three *E. coli* persisters to the nucleotide diversity of the rest of the genes on *E. coli* persisters using Wilcoxon rank-sum tests (as implemented using the Scipy module “scipy.stats.ranksums”). *P*-values were FDR corrected. Genes that had significantly higher or lower coverage (calculated using the same method) were excluded from this nucleotide diversity analysis.

### Comparative genomic analysis on *E. coli* persisters and non-persisters

Infant-specific *E. coli* persister and non-persisters genomes that were from the same subspecies clusters as the dRep-chosen *E. coli* representative genomes were used to conduct comparative genomic analysis. Identification of matching scaffolds between *E. coli* persisters and non-persisters were achieved via BLAST (Altschul et al., 1990). Specifically, scaffolds from *E. coli* persisters were compared to scaffolds from *E. coli* non-persisters using BLASTN (keeping the best hit, e-value cutoff 1e-10).

For each function that was found to be significantly enriched in *E. coli* persisters, we identified the scaffold from *E. coli* persisters in which the function was encoded on as well as the matching scaffold from *E. coli* non-persisters. Whole-scaffold alignments between persisters and non-persisters were performed in Geneious (Kearse et al., 2012). Final alignments displayed in Figure 3S were created via clinker (Gilchrist and Chooi, 2020).

### Community diversity analysis

Since the earliest fecal sample was collected several days after birth for preterm infants and around the first month of life for full-term infants, all alpha- and beta-diversity analysis between the two infant groups were conducted in the same chronological-age time frame (thus excluding any preterm samples taken before month 1). To measure convergence of the gut microbiomes, if not otherwise specified, a Wilcoxon rank-sum test was conducted to compare gut microbiomes at months 1 and 12. Modules from scikit-bio (http://scikit-bio.org/) were used to calculate the Shannon diversity index (“skbio.diversity.alpha.shannon”), weighted and unweighted UniFrac distances (“skbio.diversity.beta.weighted_unifrac” and “skbio.diversity.beta.unweighted_unifrac”, respectively), Bray-Curtis distance, and Jaccard dissimilarity (both were implemented via “skbio.diversity.beta_diversity”). To calculate UniFrac distances, a phylogenetic tree was constructed by comparing all 1005 dereplicated bacterial subspecies to each other using dRep *cluster* with a mash sketch size of 10,000.

### Principal components analysis

Principal components analysis (PCA) (performed using scikit-learn (https://scikit-learn.org/)) was conducted based on the relative abundance of bacterial subspecies in each fecal metagenome as assessed using weighted UniFrac distance. Significance of the clustering by variables (i.e., mode of delivery, prematurity, and feeding type) was determined by Permutational Multivariate Analysis of Variance (PERMANOVA) with 1000 permutations (as implemented using the scikit-bio module “skbio.stats.distance.permanova”).

## QUANTIFICATION AND STATISTICAL ANALYSIS

Statistical significance was calculated using Fisher’s exact test (as implemented using the Scipy module “scipy.stats.fisher_exact”), and Wilcoxon rank-sum test (as implemented using the Scipy module “scipy.stats.ranksums”) using the as reported in the main text and in the Methods.

### Data and Code Availability

All data that are necessary for evaluating the conclusions in the manuscript are present in the manuscript and/or the supplementary files. Software used in this manuscript is publicly available. Reads will be available under BioProject PRJNAX; SRA studies SRPX; and SRA accessions SRRX. Dereplicated *de novo* assembled genomes will be available on ggKbase (https://ggkbase.berkeley.edu/).

#### Supplemental Information

**Figure S1. Fecal sample collection of all 42 infants**

Each row represents an infant. Each solid circle corresponds to a sequenced fecal sample and they are colored by infant’s prematurity (salmon red: preterm infants; skyblue: full-term infants). Maternal fecal samples were collected around the time of delivery and are represented by solid circles before the first day of life of the matching infant. Gray boxes covered 20 infants that were removed from the downstream analysis due to well-to-well sample contamination.

**Figure S2. Some common maternal intestinal strains were transmitted and persisted in infants**

(A) Normalized relative abundances of 12 maternal bacterial strains that were detected in both samples collected two years apart. All other maternal strains were categorized as “Others” and were colored gray.

(B) Schematic of 5 of 12 maternal strains that were acquired by siblings #7 and #133. Each row represents an infant-specific, maternally transmitted strain, and strain-specific coloring is consistent with (A). Marks represent months in which the strain was detected. Shapes represent distinct strain identities (i.e., persisters, non-persisters and late colonizers). Solid marks indicate ≥99.999% popANI between the infant and mother strains, and hollow marks indicate windows in which the identity of the infant and mother strains fell below the strain cutoff. Non-persisters of the same subspecies are connected via dashed lines whereas persisters or late colonizers of the same subspecies are connected via solid lines. Double hashing indicates a non-persister early colonizer that is not maternally derived being replaced by a closely related maternal intestinal strain.

**Figure S3. Comparative genomic analysis of***E. coli* **persister and non-persister strains**

Scaffold alignments between an *E. coli* persister and an *E. coli* non-persister showing missing genetic segments including Type VI secretion system (A) and uropathogenic Escherichia coli colicin-like protein (Usp) (B) from the *E. coli* non-persister strain. Both functions were found to be associated with *E. coli* persistence in the infant gut (Tables 1 and S5). Gene alignments were generated using clinker (Gilchrist and Chooi 2020).

**Figure S4. Additional early-life community assembly dynamics of full-term and preterm gut microbiomes**

(A) Shannon indices of the gut microbiomes of full-term (left) and preterm (right) infants. Each solid circle corresponds to an infant- and month-specific Shannon index. X-axis is infant age in months and y-axis is Shannon Diversity index.

(B) Weighted (left two panels) and unweighted (right two panels) UniFrac distances comparing the gut microbiomes of full-term and preterm infants (top two panels) and infants and mothers (bottom two panels). Full-term to full-term comparisons are shown as solid sky-blue lines, preterm to preterm comparisons are shown in solid salmon-red lines, and full-term to preterm comparisons (Figure 5B,C) are shown in dashed gray lines. X-axis is infant age in months.

(C) (Top) Bray-Curtis distances comparing the gut microbiomes of preterm and full-term infants. (Bottom) Bray-Curtis distances comparing the gut microbiomes among preterm (salmon-red line) or full-term (sky-blue line) infants.

(D) Weighted UniFrac distance PCA plot of the year-1 infant and maternal gut microbiomes. Shapes and colors represent distinct sample types (preterm, full-term or maternal fecal samples).

**Table S1.** Infant metadata

**Table S2.** Details of metagenomics sequencing per fecal sample

**Table S3.** CAZymes that were significantly enriched in *Bifidobacterium* (A) and *E. coli* (B) persisters

**Table S4.** Annotations (KOs, Pfams, TCs) with corresponding genes that were significantly enriched in *E. coli* persisters

