## Supplementary figures and images for "Infant gut strain persistence is associated with maternal origin, phylogeny, and functional potential including surface adhesion and iron acquisition"

### Supplemental Figure 1

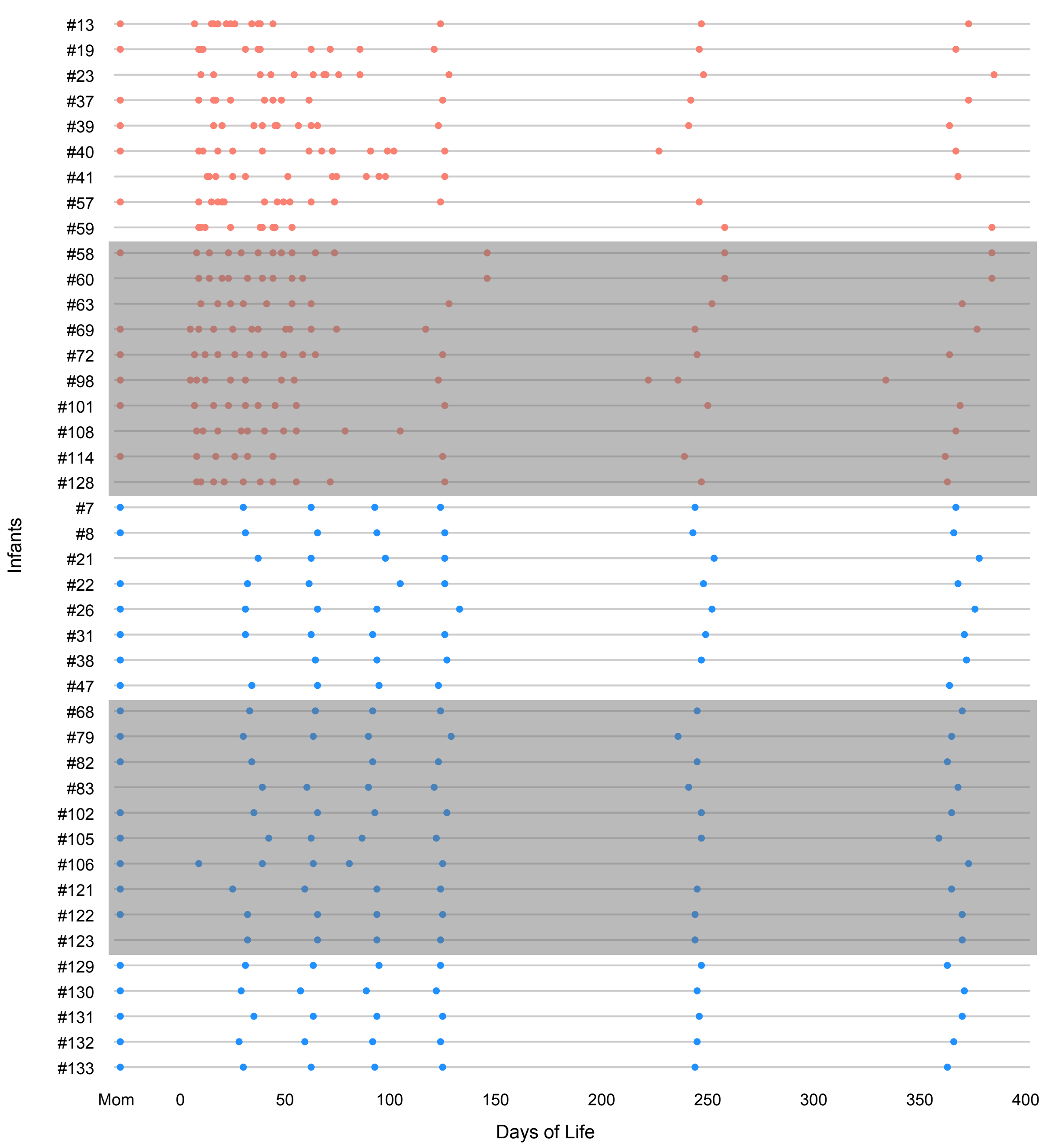

### Supplemental Figure 2

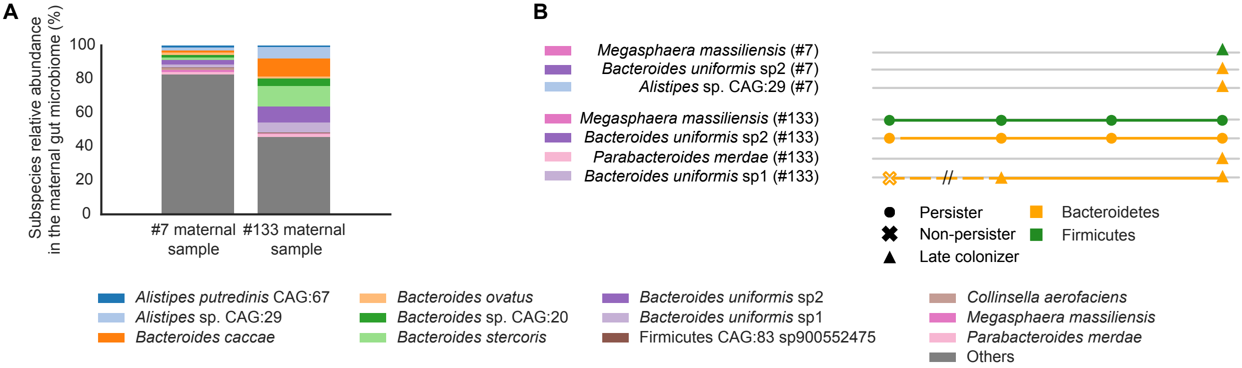

### Supplemental Figure 3

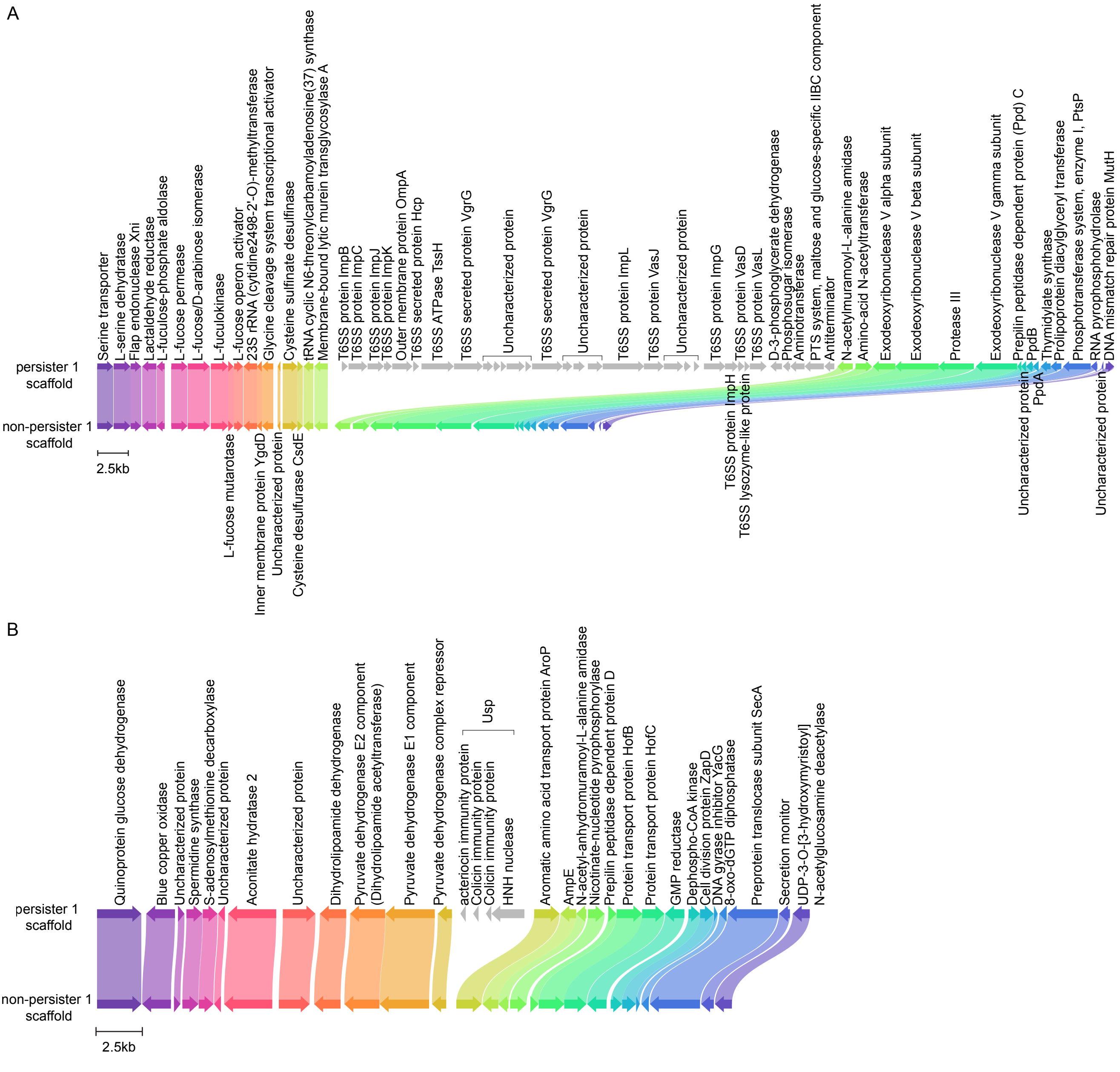

### Supplemental Figure 4

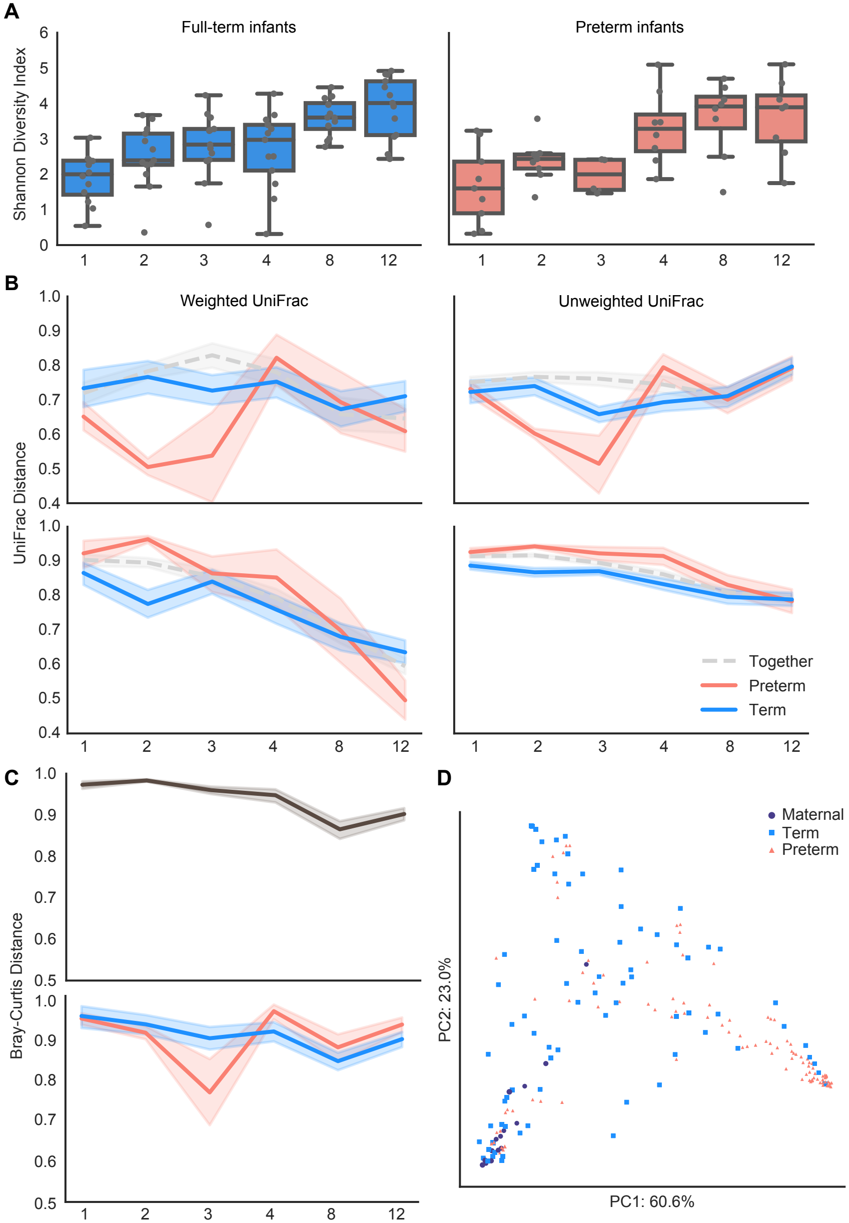
